# Analysis of fMRI data using noise-diffusion network models: a new covariance-coding perspective

**DOI:** 10.1101/223784

**Authors:** Matthieu Gilson

## Abstract

Since the middle of the 1990s, studies of resting-state fMRI/BOLD data have explored the correlation patterns of activity across the whole brain, which is referred to as functional connectivity (FC). Among the many methods that have been developed to interpret FC, a recently proposed model-based approach describes the propagation of fluctuating BOLD activity within the recurrently connected brain network by inferring the effective connectivity (EC). In this model, EC quantifies the strengths of directional interactions between brain regions, viewed from the proxy of BOLD activity. In addition, the tuning procedure for the model provides estimates for the local variability (input variances) to explain how the observed FC is generated. Generalizing, the network dynamics can be studied in the context of an input-output mapping - determined by EC - for the second-order statistics of fluctuating nodal activities. The present paper focuses on the following detection paradigm: observing output covariances, how discriminative is the (estimated) network model with respect to various input covariance patterns? An application with the model fitted to experimental fMRI data - movie viewing versus resting state - illustrates that changes in excitability and changes in brain coordination go hand in hand.

## 1 Retrospective on neuroimaging data analysis

Information processing in the brain relies on detailed interactions between many specialized neuronal subsystems that are organized in a distributed manner. The study of brain function relies on neuroimaging techniques that indirectly measure brain activity, such as electroencephalogram (EGG), magnetoencephalo-gram (MEG) and functional magnetic resonance (fMRI). In the case of fMRI, scanners record the blood-oxygen-level dependent (BOLD) signals, which are related to the energy consumption of brain cells. Early studies of task-evoked activity focused on the differences in the mean recorded BOLD level between several conditions [2]: this node-centric viewpoint allowed for tying functions (e.g., visual or auditory processing, memory, attention) to specific regions of interest, or ROIs [14]. However, in order to uncover how the brain performs tasks, the key is to understand the *coordination* between ROIs, not only which ROIs become activated. For example, the recognition and manipulation of objects in a natural environment requires the binding of visual and motor percepts together with memory.

In the 1990s, the activity of the human brain at rest has raised interest [3], because the observed large BOLD fluctuations had a structured baseline that could not be simply explained by observation noise [48]. This led to the concept of functional connectivity (FC), which describes the correlation pattern of BOLD signals for a network of ROIs [21]; see Fig. 1A for an schematic illustration with 4 ROIs. FC-based analysis has then been applied to the study of changes in the coordination between cortical regions for various tasks, for instance involving memory [50]. In this network-oriented approach, the level of correlated activity between ROIs is hypothesized to reflect the interaction strength between them, although the precise relationship between BOLD signals to neuronal activity remains only partially understood [34, 20, 32]. In comparison, for brain rhythms observed using MEG/EEG whose link with neuronal activity is clearer, synchrony-based mechanisms have been proposed to implement neural communication in a more concrete fashion [57, 6, 23].

**Figure 1:**
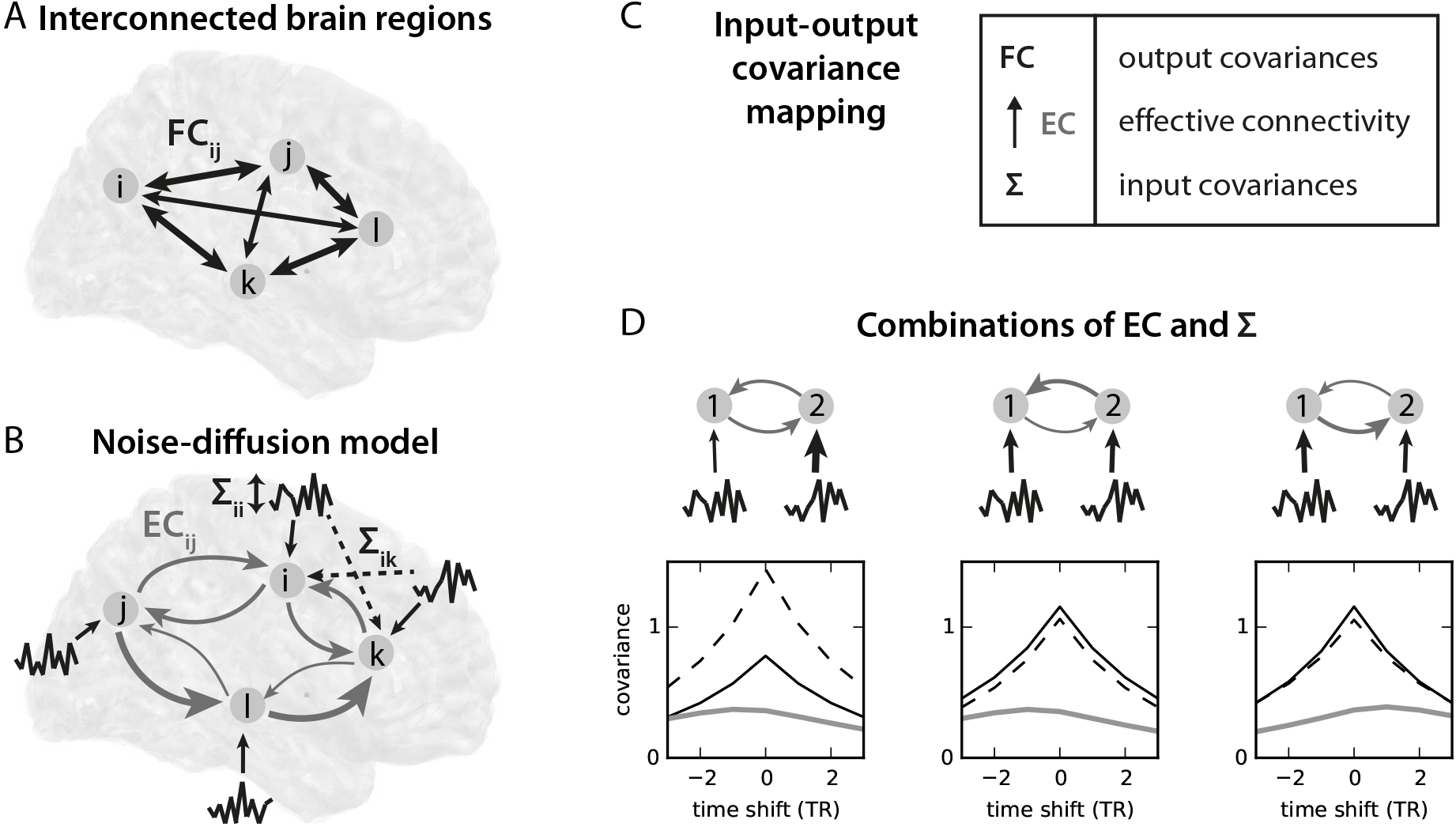
**A:** Classical fMRI analysis relies on the level of correlation between the BOLD activities of ROIs (here four), as measuring by the functional connectivity (FC) corresponding to the bidirectional arrows. **B:** The dynamic model divides the brain in a number of regions and describes the causal interactions between them, or effective connectivity (EC represented by the light gray curved arrows). Here the ROIs receive fluctuating inputs determined by their covariance matrix Σ; off-diagonal elements of Σ correspond to cross-correlated inputs. **C:** The model FC is the result of the operation of the recurrent EC on the the input covariances Σ. **D:** Three examples for the combination of EC and Σ with two connected nodes. The thickness of the gray and black arrows indicates the respective strengths for EC and Σ; note the slightly unbalanced in Σ for the right panel, which has opposite connectivity compared to the middle panel. The resulting spatiotemporal FC is displayed below each network diagram: the autocovariances of node 1 and 2 correspond to the solid black and dashed curve; the cross-covariance is displayed in thick gray. The x-axis correspond to the time shift (in TRs, the time resolution of fMRI, typically 2 s); a peak for positive time shifts indicates that 1 leads 2. The curves are calculated using Eqs. (4) and (5).

To move beyond a phenomenological description of the observed brain activity, models have been proposed to describe how regional mechanisms and network connectivity interplay to generate BOLD signals. The widely-used method of dynamic causal modeling (DCM) targets the communication between ROIs by estimating causal interactions between them, which is referred to as effective connectivity, or EC [26, 58]. In a network, EC usually shapes FC in a nontrivial fashion: weakly connected ROIs can exhibit strong correlation due to a strong global feedback [47]. For the estimation procedure, defining observables - namely how to measure the empirical brain activity and its counterpart in models - is clearly important here. Early DCM modeling aimed to reproduce BOLD time courses, e.g., succession of word-listening and silence periods [25]; recently, it has been modified to the statistics of the BOLD fluctuations measured by the cross spectra, which is the Fourier transform of covariances between ROIs [27]. Note that, for the same observables, the estimated EC values depend on the model details for the local dynamics.

Recent studies showed that the resting-state BOLD activity has a specific temporal structure, which is modulated depending on the task performed by the subject [33]. Moreover, the whole brain network exhibits a time-lag structure where some ROIs lead others at the scale of the fMRI time resolution (TR, typically 2 seconds) [43], which is also affected by behavior [44]. Note that the timescale here is different from that used for ‘dynamic FC’, of the order of a minute [9, 36]. It is important to Identify which timescales (or frequency ranges) in the BOLD signals convey information about the behavioral conditions in order to define the FC that a model should reproduce. The above-mentioned results suggest that FC should be spatio-*temporal*, not simply spatial.

In a complementary line of research, it has been argued that the study of brain function requires modeling the *whole* brain activity, in order to capture the distributed interactions across brain regions [15]. This differs from previous studies where a few ROIs were selected a-priori [30, 58], leaving many unknowns for unobserved parts of the brain. This is supported by a recent fMRI study, which demonstrated that no localized subset of cortical or subcortical ROIs could be isolated to predict the level of psychological pain rated by subjects [10]. Rather, the relevant information turned out to be globally scattered over the whole brain. Here a key point is understanding the interaction between the local dynamics and network connectivity that generates the global activity pattern [7, 18]. To investigate whole-brain communication, a specific focus of modeling studies has been on bridging bottom-up and top-down approaches by integrating anatomical and functional data [35, 41, 51]. Long-range white-matter anatomical connections are the backbone of interactions between distant brain regions and are known as structural connectivity, or SC [54]. SC can be estimated using MRI and techniques like diffusion-tensor imaging (DTI) or axonal tracing to estimate the density of the synaptic pathways connecting ROIs [13, 37, 52]. The properties of the so-called brain connectome as a graph have been extensively studied [31, 60] and are important to take into account to understand brain function [59] and when designing whole-brain models [15, 16].

## 2 Noise-diffusion network model to interpret the fMRI spatio-temporal structure

This section presents some details of a recent model [28], which was proposed to address several issues raised in the previous introductory section. This generative model aims to reproduce the BOLD activity of the whole brain (or cortex) parcellated into about a hundred ROIs [56, 31]. As depicted in Fig. 1B for 4 ROIs, the model comprises two sets of parameters:

- the spontaneous activity or local excitability received by each ROI, as described by the (co)variance matrix Σ;
- the EC strengths between ROIs, corresponding to the directional curved arrows.

Note that the framework can be extended to include input cross-covariances between ROIs corresponding to off-diagonal elements in the matrix Σ [29]; see dashed arrows in Fig. 1B. Earlier whole-brain modeling studies [15, 41] focused on adjusting the local dynamics for ROIs while using SC values for EC, which is then symmetric. In contrast, the present model is equipped with an optimization procedure to obtain an estimated EC strength for each directional connection. The rationale is that EC reflects the net effect of various mechanisms in generating FC, such as heterogeneous concentrations or types of neurotransmitters and synaptic receptors, beyond simply the density of synaptic connections that is measured in SC. SC is nevertheless used to determine the skeleton of EC (i.e., which weights are optimized and which are kept equal to 0). To obtain directional EC, the key lies in using the *spatio-temporal* FC, which captures similar information to the time-lag structure between ROIs [43, 44] and extends previous studies relying on zero-time-lag statistics of the BOLD signals - or spatial FC [8, 16, 41]. In practice, we employ the BOLD covariances without and with time shift Δ for all pairs of ROIs indexed by *i* and *j*:

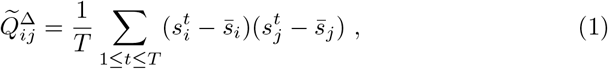

where 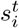 is the observed BOLD time series for ROI i. The entrainment between ROIs also depends on the local activity regime of the target for each connection. To take this into account, the model optimization also tunes the input (co)variances - the matrix Σ in Fig. 1B - in addition to EC. The optimization procedure of the whole-brain EC model [28] aims to solve the trade-off between the network size and robustness of the estimates:

- The model relies on simpler local dynamics than, e.g., the DCM [26].
- The network topology is determined by DTI measurements, which reduces the number of parameters to estimate (typically, EC has a density of 30% compared to all possible connections, giving a few thousands EC parameters to estimate for 100 ROIs).
- The inferred EC corresponds to a maximum-likelihood estimate and can be extracted from single fMRI sessions (with a duration of 5–10 min).

Formally, the vector y of BOLD activities for all ROIs obeys a multivariate Ornstein-Uhlenbeck (MOU) process:

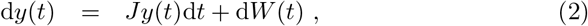

where the Jacobian *J* is defined by the EC weights for its off-diagonal matrix elements, whereas its diagonal elements are –1/*τ* for the time constant *τ* that governs a homogeneous exponential decay of the activity for all nodes; *τ* is estimated from the empirical data [28, Fig. 7B and C]. Formally, *W* is a Wiener process, corresponding to spatially correlated and temporally uncorrelated white noise. Note that the notation is adapted from the original papers. The model FC is defined as the covariances of the BOLD time series, with a time shift Δ as in Eq. (1):

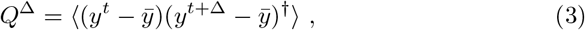

where the † superscript indicates the matrix transpose. Assuming stationarity, the following consistency equations give the model FC, defined as the second-order statistics of the nodal activities [39]:

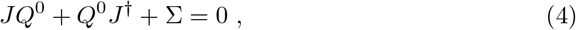

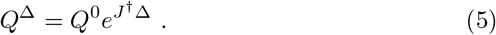

Here the covariance matrix of *ζ* is Σ = 〈*ζ^t^ζ*^†^〉, without time shift. Note that Eq. (4) implements a mapping between the input and output covariances, as schematized in Fig. 1C. The viewpoint taken on the MOU process corresponds to a noise-diffusion network, where the intrinsic variability of each node (described by Σ) propagates in the network via EC. This should be conceptually distinguished from the classical approach for linear regression, which considers that observation noise corrupts the deterministically evolving signals. In other words, distinct profiles of local variability for the nodes indicate different conditions in our context.

The combination of Σ and *J* uniquely determines the matrices *Q*^0^ and *Q*^Δ^ in Eqs. (4) and (5): this bijection between the parameters and observables allows for an unambiguous estimation [39] - up to the measurement noise of course. To reproduce the empirically observed FC, the optimization procedure [28] iteratively tunes both EC and Σ such that the model FC in Eqs. (4) and (5) best resembles its empirical counterpart in Eq. (1). It is equivalent to a natural gradient descent in the recurrently connected network with the spatio-temporal FC as an objective function [11]. To some extent, the optimization resembles a continuous-time version of Granger causality analysis of the BOLD signals [30], but it estimates the connection strengths (not their likelihood) and incorporates topological constraints given by SC.

The advantage of a proper model inversion over more phenomenological approaches [12, 43] lies in disentangling distinct contributions from Σ and EC that shape the observed BOLD statistics. For the class of linear-feedback models considered here, the observation of the lag of cross-covariances (i.e., partial observation of the temporal FC structure) or the zero-time lag covariances (spatial FC structure) is not sufficient to unambiguously estimate EC and Σ, as shown by the comparison of three simulated configurations with two nodes in Fig. 1D. The left and middle network have very similar cross-covariances (thick gray curve) with a maximum for a time shift of –1 TR. However, the origin of this observed asymmetry in the temporal FC has a different origin for the two networks: it arises from an imbalance between the Σ received by the two nodes for the left panel, whereas a stronger EC from 2 to 1 than 1 to 2 is the cause for the middle panel. Similarly, the middle and right networks have almost exactly the same values for the three curves at the origin (time shift equal to 0 TR). However, they have a very different temporal structure, each reflecting the asymmetry in EC: stronger from 2 to 1 for the middle panel, and the converse for the right panel (note that the input variances in the right panel are slightly imbalanced to obtain similar zero-lag autocovariances to the middle panel). As is well-known for processes with linear feedback, spatial information about the covariances is not sufficient to estimate the input and connectivity parameters [39]. This motivates the revised definition of (both empirical and model) FC based on covariances with time shifts.

In line with many previous definitions of EC [26, 22, 42, 4], the model FC is generated by the interplay between the network connectivity and local dynamical variables, which must be taken care of in the estimation procedure. Note that the noise-diffusion model ignores the hemodynamic response in the generation of the BOLD signals that is explicitly modeled in DCM [5, 24]. In this sense, the proposed model-based approach is more abstract and phenomenological than, for example, recent studies [1, 46] that aim to provide a mechanistic implementation of communication by synchrony/coherence for EEG and MEG data [57, 23]. Nevertheless, the parameters of the model fitted to empirical FC provide a biomarker for the brain dynamics - measured via the proxy of the BOLD dynamics - in a space of a few thousands dimensions. Such biomarkers can then be used to compare the coordination between ROIs across conditions and interpret the change thereof [29], or examine individual differences in BOLD activity [45].

## 3 Variance-based versus rate-based coding

Moving a step further to the abstract, let us consider a variant of the dynamics defined in Eq. (2) by replacing *ζ* with a time-dependent variable *x* that is characterized by its mean and its variance (discarding higher orders):

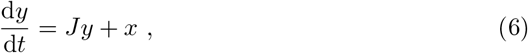

for which the linear feedback is described by the Jacobian *J*. This model defines two linear mappings, one for the means and one for the zero-lag covariances:

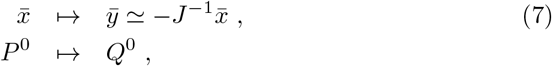

where 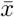 is the covariance of *x*, defined similarly to *Q*^0^ in Eq. (3) and corresponding to Σ for *ζ*. Note that the mapping for covariances depends on the input details and may differ from the Eq. (4) that is specific to the Wiener process.

These mappings can be used for detection, as illustrated for a single node in Fig. 2. The corresponding paradigm for rate detection consists in inferring changes in the input mean 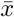 from the observation of the output mean *ȳ*, as shown in Fig. 2A. It shows the desired situation when the changes in 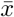 are amplified in *ȳ*, which shifts the corresponding activity histogram upwards in the right panel. As mentioned earlier, traditional approaches for task-evoked fMRI activity compare the mean activity level of ROIs between task and rest, which comply with the classical paradigm from information theory: a noisy input is characterized by its mean and the variability is considered to be observation noise or irrelevant information. A large variability for y reduces the detection accuracy by increasing the overlap between the two histograms in the right panel: for observations over short time windows, the evaluated means may fluctuate dramatically. In Fig. 2B, the input variance *P*^0^ is modified instead of the mean 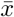, which can be detected via the output variance *Q*^0^. The second-order statistics are the basis for information here; note that the corresponding observation noise would be of higher order. In the case of fMRI, this corresponds to changes in FC. Last, when mixed effects occur as in Fig. 2C, both methods may be used to detect changes in *x*, but one may be more accurate than the other. It is not clear for fMRI data when first- or second-order statistics are more informative.

**Figure 2:**
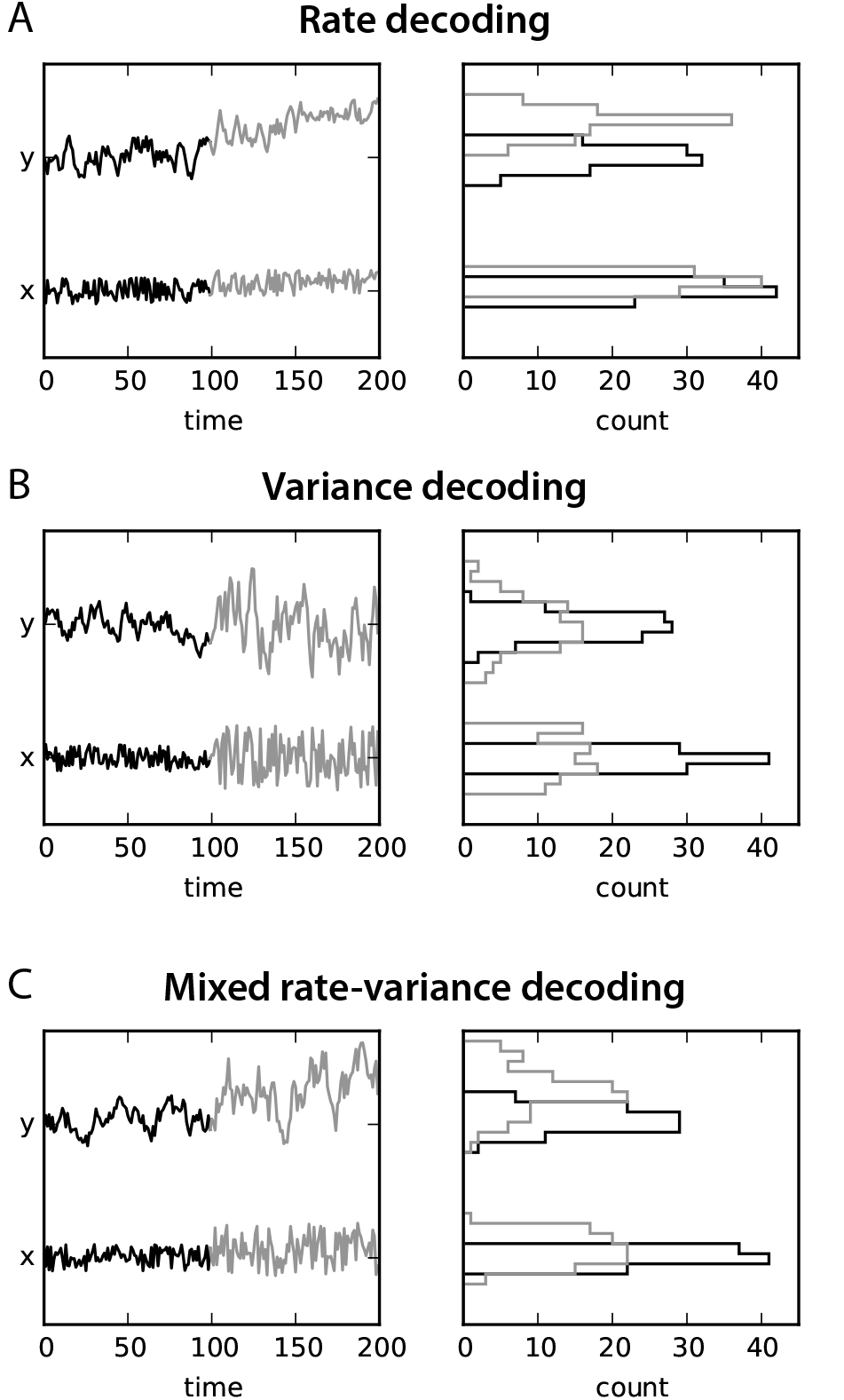
**A:** Rate-coding paradigm: a (small) change in the mean 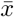 of the noisy input *x* results in a (large) change in the mean *ȳ* for *y*; each period is indicated by its color, black and gray. The quality for detection is illustrated by the distinct means for the two activity histograms in the right panel. **B:** Variance-coding paradigm: the same dynamics can be used to detect a change in the variance of *x* by observing the variance of *y*. Here, the quality of the detection relates to the distinct widths of the histograms (same color coding as in A). C: Mixed paradigm: both the mean and variance of *x* are changed between the black and gray periods, which is reflected in the histograms for *y*.

The detection scheme in Fig. 2 can be extended in two directions:

- non-zero time shifts for covariances in Eq. (7), involving *P*^Δ^ and *Q*^Δ^ for some Δ > 0;
- non-linear dynamics that dynamically regulate the gain for each node depending on the input level.

Such an non-linear model implements a rich input-output mapping for covariances, which can be used similarly to the rate mapping for information transmission. It is worth noting that the space for covariances is higher-dimensional than that for means. This is reminiscent of the “correlation theory of brain function” proposed by von den Malsburg [40]: elaborate representations of complex objects require flexible dynamics and interactions between spiking neurons. Therefore, we use the terminology (co)variance-based coding to describe the second mapping in Eq. (7). In the following, however, we focus on condition-specific EC estimated over a recording period during which the network dynamics is assumed to be stationary (in the same behavioral condition). Under these conditions, a linearized EC can be evaluated to describe the net effect resulting from the interplay between the local non-linearity and the network connectivity [16]. We leave further developments for later study.

## 4 Selectivity of network connectivity to patterns of local excitability

For fMRI, the parallel to Fig. 2B thus corresponds to flexible context-dependent coordination between brain regions (FC repertoire), which is determined by a specific EC for each behavioral condition. If this hypothesis is true, EC should be “set” in accordance to Σ for each condition, such as to constrain the propagation of local fluctuating activity (e.g., coming from sensory ROIs) to specific pathways that organize the brain coordination. Now we look into real data to see whether this phenomenon can be observed. We reanalyze the model estimates obtained from 19 subjects in two conditions: watching a movie and at rest [29]; data are available at https://github.com/MatthieuGilson/EC_estimation. Using the whole-brain dynamic model fitted to the fMRI measurements (cf. Sec. 2), we examine to which extent the model FC reflects the changes in the estimated local excitability Σ for the two conditions. Changes in the estimated Σ presumably relate to the stimulus load; note that Σ is diagonal here, contrary to [29].

First, we verify in Fig. 3A that the two sets of model FCs (over all subjects) reflect the two conditions. To do so, we compare the matrix distances between FC from all subjects in the same condition (gray arrow in the left panel) or between the two (black arrow). Cross-covariances are discriminative against the two conditions, as indicated by the significantly larger values for the inter-condition distances in the right panel. In contrast, the distance distribution over the subjects for variances in the movie condition is similar to the distance distribution between FCs in the two conditions (middle panel).

**Figure 3:**
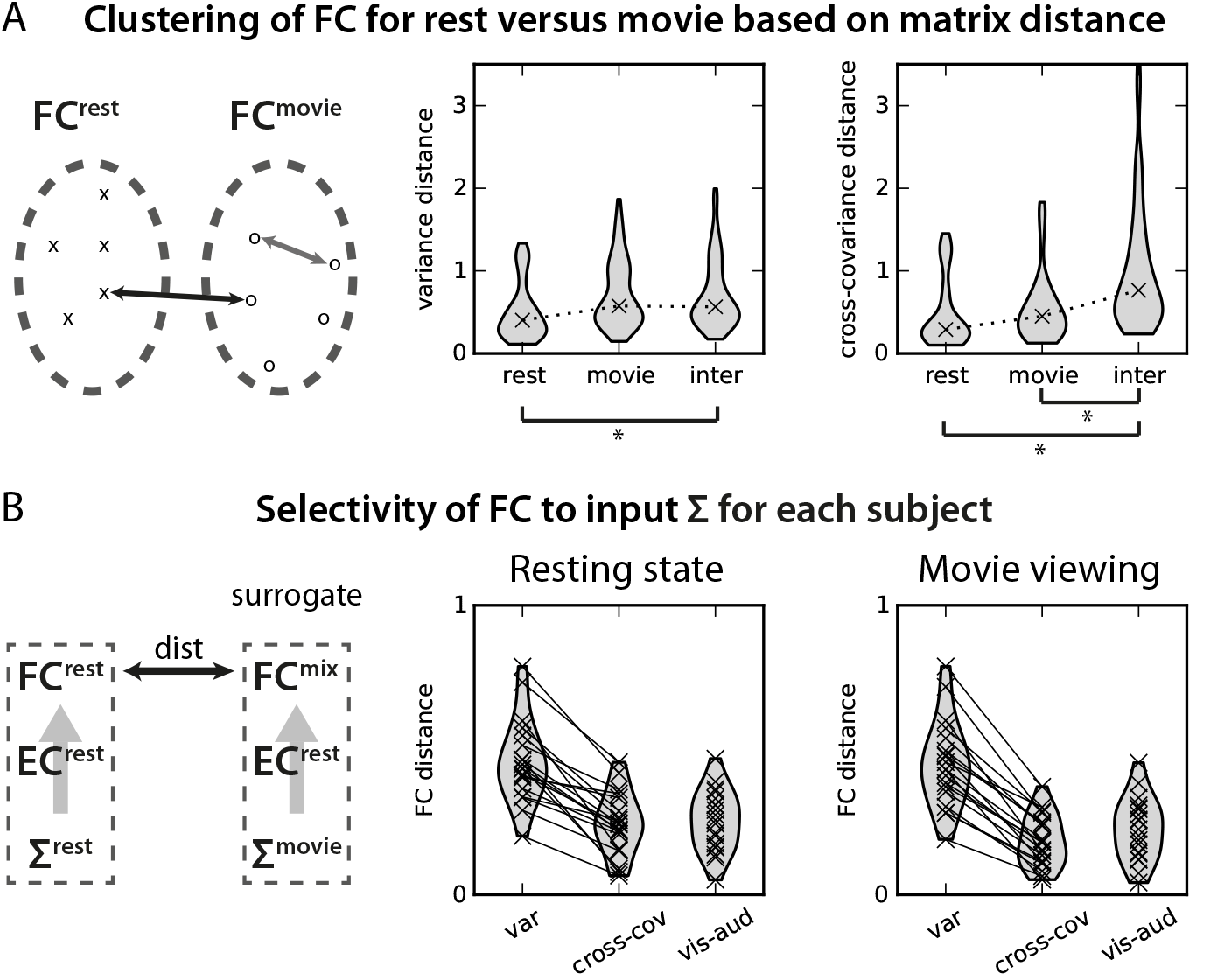
**A:** Comparison of the model FCs for the model fitted to empirical data [29]: 19 subjects were scanned in two conditions, at rest and when watching a movie. As shown in the left diagram, the FC distances are calculated for all subjects within the same condition (same symbol in the left diagram, as indicated by the gray arrow) and between the two conditions (distinct symbols indicated by the black arrow and corresponding to ‘inter’ in the middle and right plots). The middle plot displays the distances over all pairs of subjects for the variances (the medians are indicated by the crosses), the right plot for the cross-covariances; note that this distance measure is normalized by the mean empirical FC averaged over the two conditions and for the corresponding matrix elements. Significant differences between the ‘cross’ distribution and rest/movie are indicated by the stars below (*p* < 0.01 for Welch’s t-test and Mann-Whitney test); stars for the comparisons between rest and movie are not shown, although they are significant. **B:** To quantify the selectivity of FC with respect to Σ for each subject, the model FC fitted to the empirical data - corresponding to EC and Σ for the same condition (here rest) - is compared with a surrogate mixed FC when swapping Σ for the other condition, as illustrated in the left diagram; only the spatial FC corresponding to *Q*^0^ in Eq. (4) is used here. Here the distances are normalized to give 1 for a matrix compared to the null matrix with zeros everywhere. The middle plot shows the distribution of normalized matrix distances (as in A) between FC rest and mix for EC in the rest condition (each cross representing a subject), and the right plot for EC in the movie condition. The distance between the two FCs is calculated for several subsets of the matrix elements: variances on the diagonal, cross-covariances outside the diagonal and 14 visual/auditory ROIs.

Second, Fig. 3B examines to which extent the FC variances and crosscovariances (diagonal and off-diagonal matrix elements, respectively) are the expression of the condition-specific Σs. The corresponding model FCs obtained for the same EC, but distinct Σ (see left diagram) are compared using a normalized matrix distance; moreover, this is decomposed for the matrix elements indicated on the x-axis of the middle and right panels in order to quantify their relative contribution in shaping FC. Although variances are modified to a greater extent than cross-covariances by swapping the conditions for Σ, the cross-covariances turn out to be impacted as well. Interestingly, ECs that are highly selective for variances are selective for cross-covariances too, as indicated by the lines connecting the crosses for the same subject. This means that, for each subject and condition, the mapping determined by EC is not independent from the Σ it is paired with. Among cross-covariances, those concerning visual and auditory ROIs are more affected than on average (‘vis-aud’ label), which makes sense for the passive viewing and listening task. The interpretation is that these high-activity ROIs feed the brain network in the movie condition [29].

## 5 Future perspective for fMRI analysis and quantifying whole-brain communication

As mentioned in the beginning of the paper, the state of the art for the analysis of task-evoked fMRI activity has shifted from studying the means of BOLD signals to their second-order statistics, that is, from a structure-centric viewpoint (one region = one function) towards a network-oriented viewpoint, focused on network interactions and correlation pattern [27, 15, 19]. Here a complementary aspect has been highlighted based on the input-output mapping implemented by the interactions between ROIs (as quantified by EC) in a whole-brain dynamic model [28], which provides a mechanistic description of the interplay between local and network properties in the generation of FC. This model appears particularly interesting in regards of new possibilities to stimulate the brain in a non-invasive manner [53]: the modification of the local activity for certain ROIs can be evaluated with the model to explore the resulting effects at the network level. Here the discriminative power EC with respect to Σ corresponds to the efficiency of stimulation protocols in changing FC. As another example, therapies based on neurofeedback - where the patient sees her/his brain activity and attempts to manipulate it - currently rely on changes in FC as well as in BOLD activity level [38]. One can move one step further with the model and aim to shape directional (BOLD) interactions between brain regions. The proposed model with linear feedback [28] is arguably the simplest dynamics to take time into account in the generation of FC, assuming stationarity for a given condition; more detailed mechanisms should be incorporated in the dynamic model in order to deepen its interpretation capabilities. However, the multiplication of parameters is expected to reduce the estimation robustness, which is problematic to obtain condition-specific estimates.

## Acknowledgements

MG acknowledges funding from from the Marie Sklodowska-Curie Action (grant H2020-MSCA-656547) and the Human Brain Project (ramp-up phase, grant FP7-FET-ICT-604102).

The author thanks Moritz Deger and Martin Nawrot for organizing the 12th International Neural Coding Workshop, NC2016. The author is also grateful to Pierre Yger, Ruben Moreno-Bote, Vicente Pallarez, Andrea Insabato, Gustavo Deco and Morten Kringelbach for constructive discussions.

